# Microtubules are necessary for proper Reticulon localization during mitosis

**DOI:** 10.1101/516773

**Authors:** Zane J. Bergman, Ulises Diaz, Amanda Sims, Blake Riggs

## Abstract

During mitosis, the structure of the Endoplasmic Reticulum (ER) displays a dramatic reorganization and remodeling event, however the mechanism driving these changes is poorly understood. Recently, the Reticulon family of ER shaping proteins has been identified as possible factors to promote these drastic changes in ER morphology. In addition, the Reticulons and other ER shaping proteins have been directly linked to several hereditary neurodegenerative disorders. Here, we provide key insight into the cytoskeletal factors involved in the Drosophila Reticulon, Reticulon-like 1 (Rtnl1) during mitosis in the early embryo. At prometaphase, Rtnl1 localizes at the spindle poles just prior to the bulk of ER localization suggesting a role in recruitment. Using precise temporal injections of cytoskeletal inhibitors in the early syncytial Drosophila embryo, we show that microtubules, not microfilaments are necessary for proper Rtnl1 localization and function during mitosis. Lastly, we show that astral microtubules are necessary for Rtnl1 localization at the spindle poles early in mitosis. This work highlights the role of the microtubule cytoskeleton in Rtnl1 localization and ER dynamics during mitosis and sheds light on a pathway towards inheritance of this major organelle.

## Introduction

The organization and dynamics of eukaryotic cells has been extensively studied over the past several decades with key discoveries made with respect to the chromosomes, and cytoskeleton network (McIntosh *et al*., 2002; Walczak *et al*., 2010; Lancaster and Baum, 2014). However, lesser understood is the dynamics of the cytoplasmic organelles during cell division. There have been studies that have investigated organelle dynamics during mitosis, including the disassembly and reassembly of the Golgi apparatus (Altan-Bonnet *et al*., 2004; Persico *et al*., 2009) and the changes in structure and localization of the Endoplasmic Reticulum (ER) (McCullough and Lucocq 2005, Lu et al. 2009). Furthermore additional studies have shed light on the regulatory mechanism involved in these dramatic changes of the Golgi (Bisel *et al*., 2008) and the ER (Bergman *et al*., 2015). While there have been several advances of our understanding of organelle dynamics during mitosis, several unanswered questions still remain, including the role of the cytoskeletal network during mitosis, the mitotic kinases and motor proteins involved in these dramatic reorganization events.

The microtubule network experiences a dramatic reorganization event to form a bipolar spindle responsible for attaching to and partitioning the genetic material. This relies on coordination of the mitotic kinases, several microtubule associate proteins and both plus-end and minus-end directed motors. Earlier studies have shown that the ER shares a close interdependence with the microtubule network (Terasaki *et al*., 1986; Waterman-Storer and Salmon, 1998) but it is unclear if microtubules play a role in these mitotic ER reorganization events. Furthermore, an early study focusing on mitotic ER dynamics in C.elegans suggest that the actin cytoskeleton is involved in these dramatic reorganization events (Poteryaev *et al*., 2005). Imaging and functional studies focused on the ER during mitosis shows that the ER shares a close association with the mitotic spindle and spindle poles and that microtubules are important for nuclear envelope reformation, in which the ER plays an important role (Bobinnec *et al*., 2003; Anderson and Hetzer, 2008; Lu *et al*., 2009; Bergman *et al*., 2015). It is important to note, however, the factors associated with this mitotic ER reorganization are currently unknown.

Recently, a family of ER shaping proteins, the Reticulons were identified as factors responsible for the generation of curvature of the tubular ER network (Voeltz *et al*., 2006). These proteins form oligomeric complexes and work in concert with other ER shaping proteins, Spastin, Atlastin, REEPs, and DP1 (Hu *et al*., 2008; Shibata *et al*., 2008). Furthermore, defects in the function of these proteins have been linked to several neurodegenerative diseases including hereditary spastic paraplegias and Alzheimer’s disease (Sanderson *et al*., 2006; Yang and Strittmatter, 2007). The role of the Reticulon proteins in their ability to control ER structure has been well documented (Yang and Strittmatter, 2007), however there have been very limited studies that suggest these Reticulon family member also bind to microtubules (Schlaitz, 2014). Additionally, the functional role of Reticulon proteins during mitosis is currently unknown.

Once fertilized, the Drosophila embryo experiences a series of 13 rapid synchronous nuclear syncytial divisions before each of the 5,000 nuclei are encapsulated by membrane furrows to form multicellular embryo (Tram *et al*., 2002). During these syncytial divisions, the first 9 divisions occur in the interior of the embryo, after the 9^th^ division, the nuclei migrate to the cortex of the embryo and go through 3 more rounds of division. These syncytial divisions are rapid and do not have gap phases in their cell cycle, rather just experience a S-phase (DNA replication) and mitosis, where the DNA is partitioned to the daughter nuclei. It is during these cortical nuclear divisions (cycle 10 – 13) that provide an excellent platform to examine mitotic events. Additionally, several studies have demonstrated that syncytial mitosis in the early embryo is comparable to mechanism of mitotic events found in other systems (Fenton and Glover, 1993; Sharp *et al*., 2000; Brust-Mascher, 2002; Frescas *et al*., 2006). Furthermore, the ER demonstrates a dramatic rearrangement in both structure and localization during mitosis in the syncytial embryo, in line with the mitotic ER changes seen in other systems (Bobinnec *et al*., 2003; Bergman *et al*., 2015).

Here we sought to examine the necessity of key mitotic components in the reorganization of the ER during mitosis. We performed microinjections of small molecule inhibitors targeting the cytoskeleton, major mitotic kinases, and motor proteins known to play a key role in mitotic progression. Using the Drosophila Reticulon, Reticulon-like 1 (Rtnl1), we show that microtubules, not microfilaments, are necessary for reorganization of the ER during mitosis. We also demonstrate that localization of Rtnl1 at the poles relies on the mitotic kinases Aurora B and Polo. The results presented here provide an initial framework of the mechanism involved in mitotic ER reorganization and provides insight on a pathway towards ER inheritance during cell division.

## Results

### Rtnl1 localizes to the perispindle region and spindle poles during mitosis in the early Drosophila embryo

In order to investigate mitotic ER dynamics, we used the GFP labeled Drosophila Reticulon-like 1 (Rtnl1-GFP), which marks the ER and is the only functional Reticulon in Drosophila (Wakefield and Tear, 2006). We examined the localization of Rtnl1 during mitosis of the cortical syncytial divisions (cycles 10-13) in the Drosophila embryo. Here we used time-lapse confocal microscopy and imaged the localization of Rtnl1 at the different stages of mitosis. The Rtnl1-GFP tagged line was identified in a protein trap screen (Morin *et al*., 2001) with the GFP tag being in frame with the endogenous locus of the reticulon gene. We created a transgenic line expressing both Rtnl1-GFP and the DNA marker, His2Av-RFP (H2-RFP) to monitor the stages of mitosis. Rtnl1 was spread throughout the cytoplasm during interphase, however as the embryo enters mitosis, Rtnl1 began to accumulate at the nuclear envelope (Fig. 1A). At prometaphase, as the nuclear envelope destabilizes, there was a rapid localization of Rtnl1 at the poles (Fig. 1A, arrow). Rtnl1 accumulated at the poles during metaphase and also was found along the perispindle region. This localization persisted through anaphase, as the chromosomes are segregated. At telophase, Rtnl1 surrounds the decondensed chromosomes and localized to the midbody.

**Figure 1.**
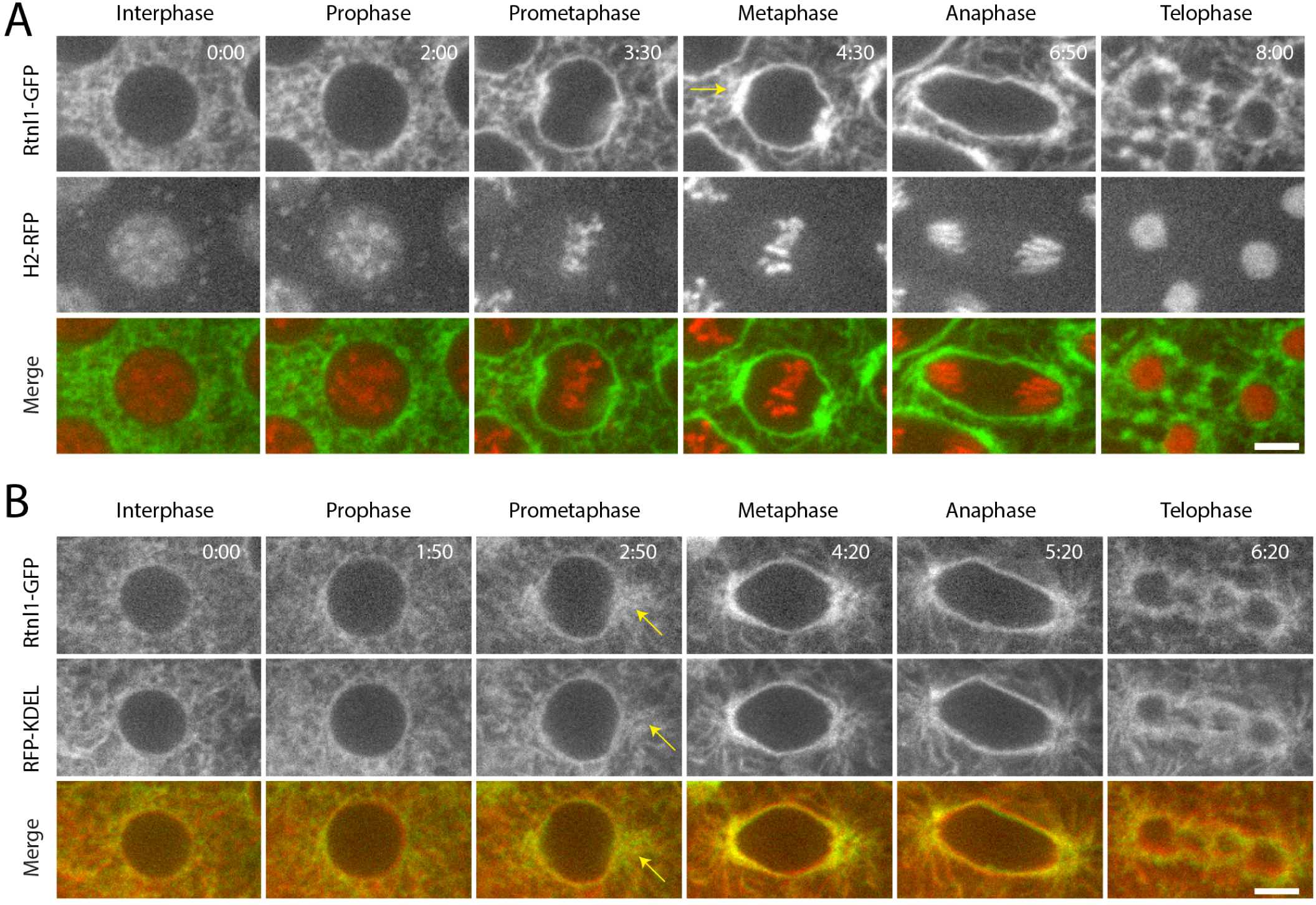
Localization of Rtnl1-GFP During Syncytial Mitoses. **(A)** Stage 10 Drosophila embryo expressing both Rtnl1-GFP (green) and H2-RFP (red). Rtnl1-GFP is spread out in a mesh network during interphase. Condensation and alignment of chromosomes (H2-RFP) shows the progression of mitosis. At onset of mitosis, an increase in GFP signal occurs at the nuclear periphery, most notably at the poles (yellow arrow). Rtnl1-GFP follows the nuclear periphery throughout metaphase and anaphase. It is then divided into the two daughter nuclei with a series of strings between them and bright staining at the midbody. **(B)** Stage 10 Drosophila embryo expressing Rtnl1-GFP (green) and RFP-KDEL (red) as it progresses through mitosis. RFP-KDEL resides within the lumen of the ER while Rtnl1-GFP is a peripheral membrane protein. These proteins overlap in much of their localization, except at prometaphase where there appears to be a bloom of Rtnl1 localization at the poles (yellow arrow). Time is in min:sec. Scale bars are 10μm.

Previous studies showed that Rtnl1 is highly colocalized with the ER during interphase (Voeltz *et al*., 2006; Wakefield and Tear, 2006), here we sought to examine the colocalization with the ER during mitosis. Transgenic embryos containing Rtnl1-GFP and a red fluorescent-tagged ER marker, RFP-KDEL were examined during the stages of mitosis (Fig 1B). Throughout mitosis, there appeared to be a high degree of localization between the ER and Rtnl1. However, as the embryo enters prometaphase, Rtnl1 appeared slightly ahead of the bulk localization of ER (Fig. 1B, arrow). Here we conclude that Rtnl1 displays dramatic changes during mitosis and is highly colocalized with the ER. Additionally, at prometaphase, Rtnl1 appears to accumulate at the spindle poles just prior to the bulk localization of ER. This result suggests that Rtnl1 is responsible for the movement of ER membrane towards the spindle poles during mitosis.

### Accumulation of Rtnl1 precedes major reorganization of the ER at the poles

Prior studies have shown that there is a strong accumulation of ER membrane at the poles beginning at prometaphase after nuclear envelope breakdown through metaphase (Bobinnec *et al*., 2003; Bergman *et al*., 2015). In addition, based on our colocalization studies (Fig. 1), we sought to follow up on our observation of Rtnl1 localization at the spindle poles at prometaphase. We closely examined the localization of Rtnl1-GFP and RFP-KDEL during the transition from prophase to prometaphase during mitosis using real-time confocal microscopy (Fig. 2A). Low magnification images showed a brief but strong localization of Rtnl1-GFP at the poles prior to localization of RFP-KDEL. High magnification insets and precise time measurements during prometaphase showed that Rtnl1-GFP localized at the spindle poles 20 seconds prior to localization of RFP-KDEL (Fig. 2B). We quantitated this localization by measurements of fluorescence intensity of Rtnl1-GFP and RFP-KDEL at the poles during prometaphase (Fig. 2B, yellow box).

**Figure 2.**
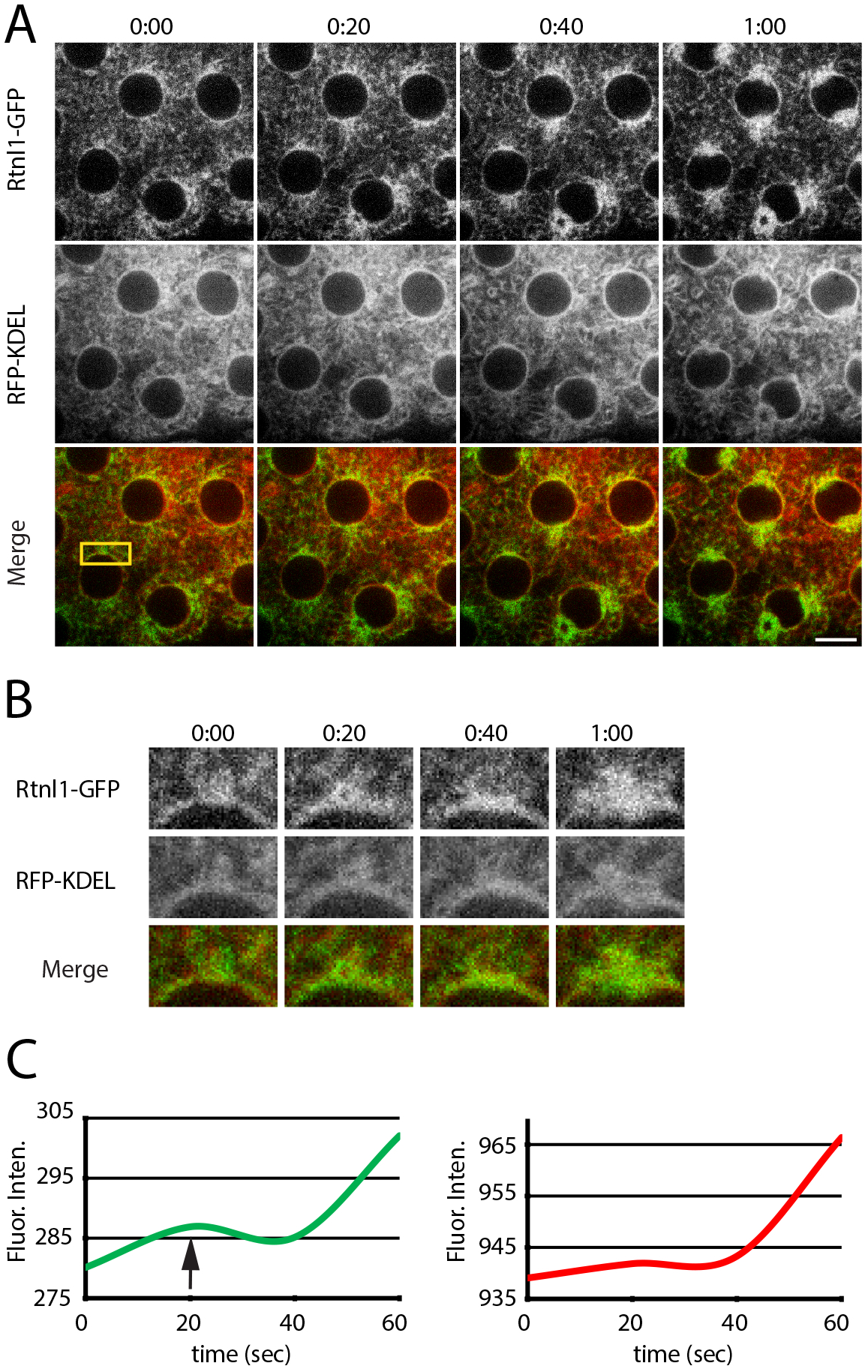
Enrichment of Rtnl1-GFP precedes major reorganization of ER to the poles early in mitosis. **(A)** Stage 10 Drosophila embryo expressing Rtnl1-GFP (green) and H2-RFP (red) as it proceeds through one minute during the early stages of mitosis. GFP signal accumulation at the poles arrives and intensifies before the RFP signal, indicating an enrichment of Rtnl1-GFP before an equal amount of ER. **(B)** High-magnification inset of one of the poles indicated by a box in A. **(C)** Plots of integrated densities of the boxes from B. There is a smaller increase of GFP signal at the 20 second time point before a rapid increase in both GFP (green trace) and RFP (red trace) at the 1 minute time point, indicating that Rtnl1-GFP does accumulate at the poles before KDEL-RFP (arrow). Time is in min:sec. Scale bar is 10μm.

Intensity measurements were taken for 1 minute at the end of prophase and at the start of prometaphase. The start and timing of prometaphase was signaled by the breakdown of the nuclear envelope and disappearance of Lamin B (Supplemental Fig 1). Measurements showed a peak of intensity of GFP at 20 seconds (Fig. 2C, arrow), while the fluorescence intensity of RFP remained low until 40 seconds into prometaphase. Taken together, we conclude that Rtnl1 localization enriches at the spindle poles during prometaphase prior to the bulk accumulation of ER membrane.

### Proper localization of Rtnl1 depends on a dynamic microtubule network

Localization of Rtnl1 at the spindle poles before bulk ER membrane localization suggest a role of microtubules for ER membrane movement during mitosis. Previous studies have shown the requirement of the microtubule network on proper ER structure during interphase (Terasaki *et al*., 1986; Klopfenstein *et al*., 1998; Waterman-Storer and Salmon, 1998), yet, little is known about the role of microtubule dynamics during mitosis. To investigate the involvement of microtubule network during mitosis, we employed precise temporal injections of small molecule microtubule poisons, just prior to the start of mitosis in the early syncytial embryo (Fig. 3B, C). Rtnl1-GFP / H2-RFP embryos were injected with the microtubule depolymerizing agent, colchicine just prior to entry into mitosis in cycle 10, after the nuclei have migrated to the cortex. Rtnl1 still was able to accumulate along the nuclear envelope, however at NEB, the ER membrane became disorganized and displayed severe clumping (Fig. 3B arrow).

**Figure 3.**
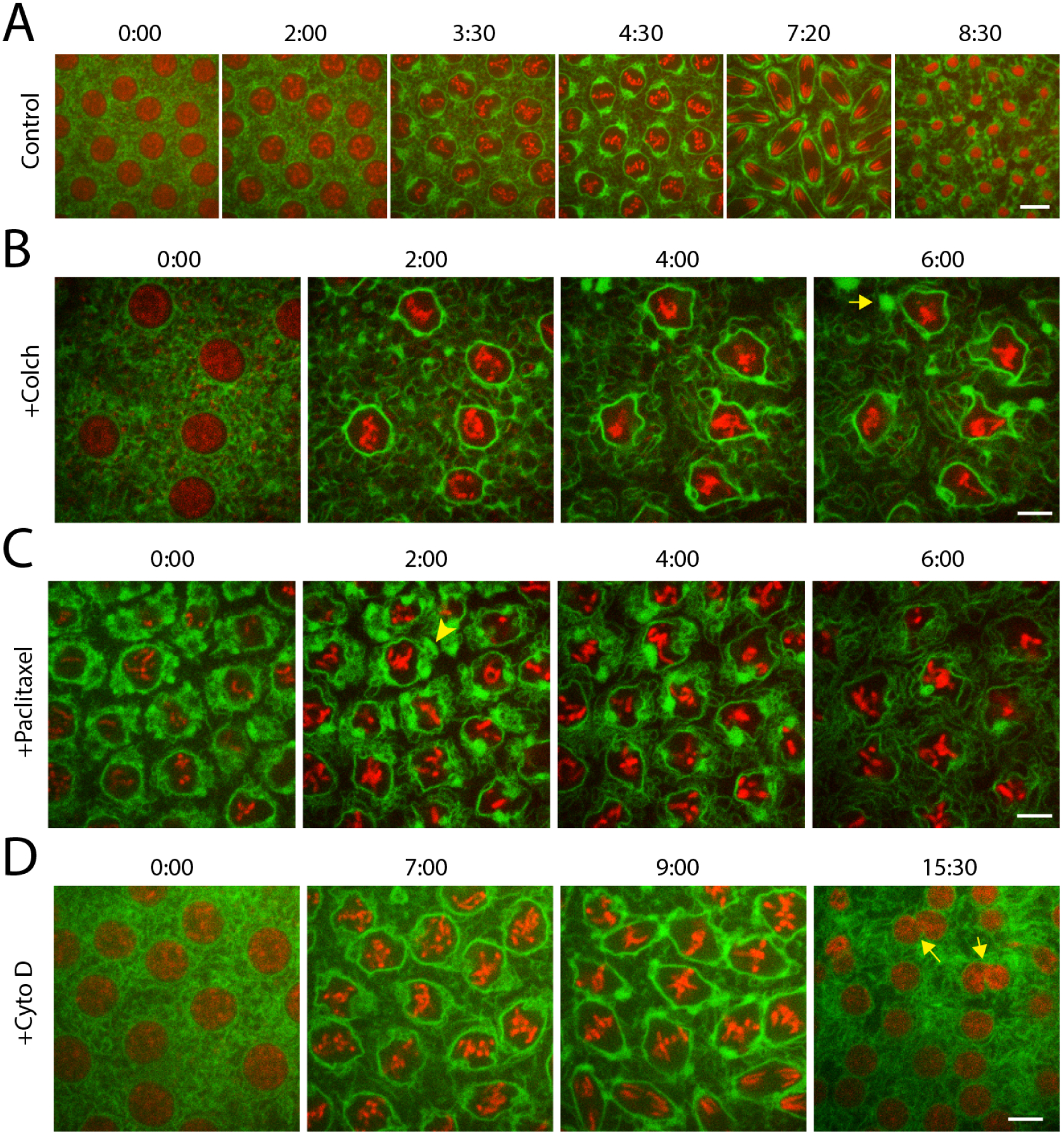
Proper localization of Rtnl1-GFP depends upon a dynamic MT network. **(A)** Control injection of a stage 10 Drosophila embryo expressing Rtnl1-GFP (green) and H2-RFP (red). Embryo was injected with 10% DMSO injection buffer and followed. No defects were observed in ER localization or chromosome condensation and alignment and there was no delay in any stages of mitosis. **(B)** Injection of colchicine into a stage 9 embryo expressing Rtnl1-GFP and H2-RFP. Though chromosomes could condense, they did not align at metaphase, arresting the embryo. Rtnl1-GFP did gather at the nuclear periphery, but did not enrich at the poles. As the arrest continued, inter-nuclear ER became tubular and the ER at the nuclear periphery lost its even distribution, snapping back to large packets (yellow arrow). **(C)** Injection of taxol into a stage 10 embryo expressing Rtnl1-GFP (green) and H2-RFP (red). Chromosomes condensed and aligned at the metaphase plate, but could not proceed, arresting the embryo. Rtnl1-GFP accumulated at the poles during the arrest (yellow arrowhead). This accumulation was then lost as the arrest continued. **(D)** Stage 9 embryo expressing Rtnl1-GFP (green) and H2-RFP (red) was injected with Cytochalasin D to disrupt actin assembly. A lack of actin cages did not affect Rtnl1-GFP organization during mitosis or chromosome condensation and movement. However, at telophase, daughter nuclei that were within proximity fused (yellow arrows), creating large nuclei. Time is in min:sec. Scale bars are 10μm.

Interestingly, the ER still remodeled into fenestrated mitotic clusters. However, proper reorganization of the ER structures at the poles is severely disrupted. Injection of the microtubule stabilizing agent Paclitaxel, prior to entry into mitosis in cycle 11 initially did not affect the early reorganization of Rtnl1 along the nuclear envelope and accumulation at the poles (Fig 3C, arrowhead). However, as mitosis progressed, Rtnl1 organization destabilized at the spindle poles and the perispindle region. Treatment of either colchicine or Paclitaxel caused a prolonged mitotic arrest at metaphase in which ER organization and structure were completely lost. It is important to note that injection of the drug vector, 10% DMSO (Fig. 3A) did not display any defects in mitotic progression and imaging of the microtubule dynamics during mitosis in the presence of either microtubule poisons in the early embryo displayed the known effects on the microtubule network. A previous study examining the remodeling and reorganization of the ER in the early C. elegans embryo suggested that the actin cytoskeleton mediated the late mitotic transitions of the ER (Poteryaev *et al*., 2005). In order to investigate the role of the actin network in mitotic Rtnl1 organization, we injected the actin polymerization inhibitor, cytochalasin D (cytoD) just prior to entry into mitosis in cycle 11. Rtnl1 localization and ER distribution appeared normal with Rtnl1 localizing at the spindle poles and along the perispindle region. It is important to note that treatment with cytoD did not cause a mitotic arrest. These embryos did display severe defects in nuclear spacing as previously published (Field and Alberts, 1995; Riggs *et al*., 2003), which led to a few cases of nuclear fusion (arrow in 3D). These results indicate that microtubule dynamics are necessary for proper reorganization of Rtnl1 at the spindle poles and along the perispindle region during mitosis and this mitotic reorganization is independent of the actin cytoskeleton.

### Accumulation of Rtnl1 at the spindle poles early in mitosis relies on a dynamic astral microtubule network

We next wanted to investigate the role of astral microtubules in recruitment of Rtnl1 at the spindle poles upon entry into mitosis. There are several Drosophila mutations that affect centrosome function, however many of these are required maternally; exhibit gross defects in nuclear spacing, chromosome segregation and spindle assembly and fail to develop (Gonzalez *et al*., 1998; Megraw *et al*., 1999). In order to examine the role of astral microtubules and to avoid these confounding effects, we used precise temporal injection of small molecules that have been shown to target astral microtubule populations. Here we injected the Aurora B kinase inhibitor, Binucleine 2 into Rtnl1-GFP / H2-RFP transgenic embryos just prior to entry into mitosis in cycle 10. Aurora B is a well-known regulatory kinase that targets many events in mitosis including chromosome condensation and transmission (Giet and Glover, 2001; Resnick *et al*., 2006) and the small molecule Binucleine 2 has been shown to specifically inhibit Aurora B isoforms in Drosophila (Smurnyy *et al*., 2011). In addition, several studies have also linked Aurora B to spindle dynamics including phosphorylation and inactivation of MCAK, a kinesin 13 family member that destabilizes microtubules (Kwon *et al*., 2004; Lan *et al*., 2004). Binucleine 2 was injected just prior to entry into mitosis and at prophase, reorganization and accumulation of Rtnl1 at the nuclear envelope appeared normal, however there was a lack of Rtnl1 accumulation at the spindle poles (Fig. 4A arrow). During the metaphase to anaphase transition, there was a minor accumulation of Rtnl1 at the poles (Fig. 4A 10:00 time point), but much less than controls (see Fig1A). There were also defects seen at telophase with lagging chromosomes present, and a lack of localization of Rtnl1 at the midbody (Fig 4A, 12:30 time point). The lagging chromosome phenotype and midbody defects are consistent with well established roles of Aurora B in chromosome alignment and central spindle / midbody formation (Ditchfield *et al*., 2003; Gruneberg *et al*., 2004; Bastos *et al*., 2013). In addition, inhibition of Aurora B dramatically affects microtubule dynamics by the continued activation of MCAK, thereby promoting depolymerization of microtubule network. These results indicate that localization of Rtnl1 at the spindle poles relies on microtubule dynamics.

**Figure 4.**
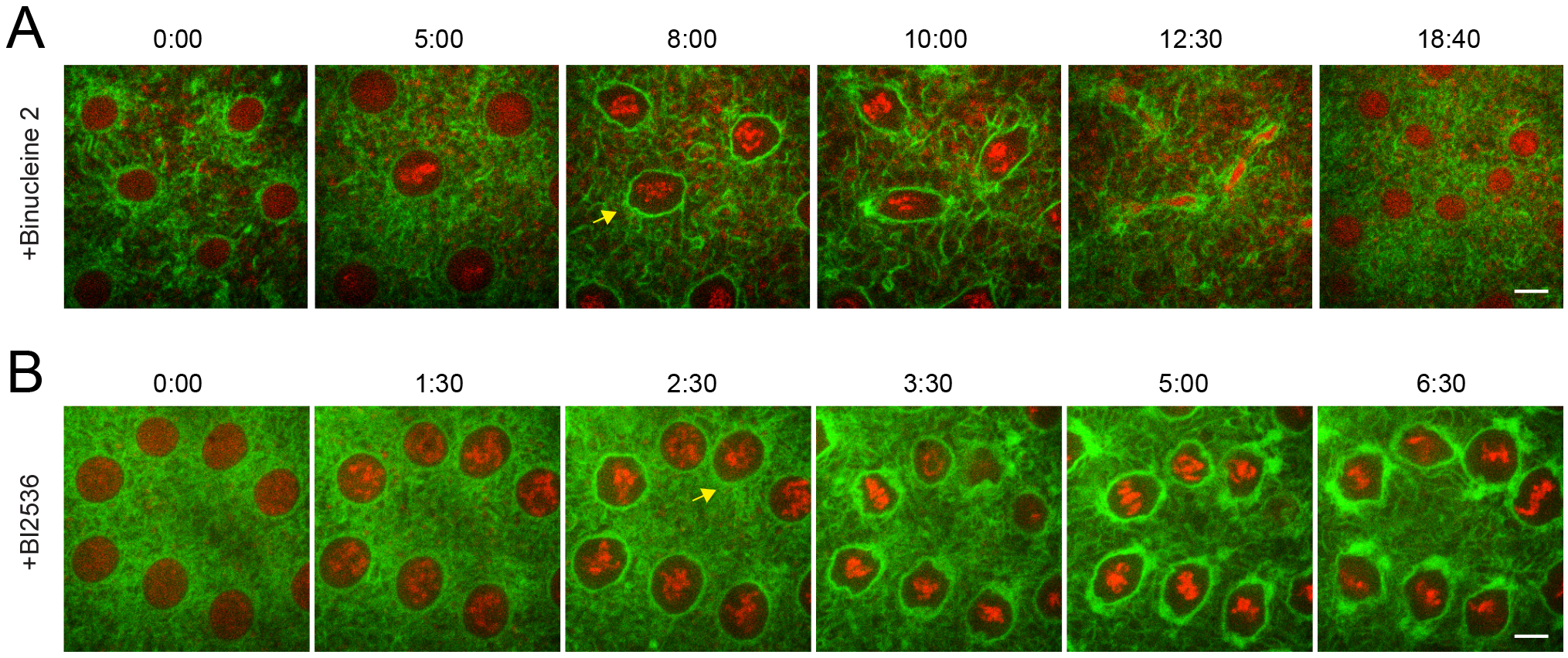
Rtnl1-GFP reorganization is dependent upon cell cycle kinases. **(A)** Stage 9 Drosophila embryo expressing Rtnl1-GFP (green) and H2-RFP (red) was injected with the aurora B kinase inhibitor, Binucleine 2. Although the embryo proceeded through the next mitosis with a delay, several defects occurred. There was a lack of GFP accumulation at the poles until anaphase (yellow arrow). Telophase was also disturbed. **(B)** Injection of a stage 9 embryo expressing Rtnl1-GFP (green) and H2-RFP (red) with BI2536, which inhibits Polo kinase. This compound eventually arrested the embryo at a metaphase-like stage. Again, there was a lack of GFP accumulation before metaphase (yellow arrow). There was a greater than normal accumulation of Rtnl1-GFP at the nuclear periphery. Time is in min:sec. Scale bars are 10μm.

In order to specifically examine astral microtubule populations, we injected the Plk1 inhibitor, BI25356 in Rtnl1-GFP / H2-RFP embryos just prior to entry into mitosis in cycle 10. The highly conserved kinase, Polo like kinase 1 (Plk1) regulates many aspects of mitosis including centrosome maturation and migration, cohesion dissociation, and chromosome segregation (Van Vugt and Medema, 2005). Importantly, a recent study also showed that inhibition of Plk1 by BI25356 specifically targets astral microtubule populations early in mitosis prior to metaphase (Serio *et al*., 2011). Rtnl1 displayed a strong accumulation to the nuclear envelope at prophase, however there was a lack of accumulation at the centrosome at prometaphase (Fig. 4B, arrow). As the embryo progressed into metaphase, there was a gradual accumulation of Rtnl1 at the poles. The embryo arrested at metaphase and there was a strong localization of Rtnl1 at the poles and along the perispindle region. Previous studies involving the Plk1 inhibitor, BI25356 displayed defects in centrosome migration and separation, astral microtubule formation at prophase and activation of the spindle assembly checkpoint (Lénárt *et al*., 2007; Serio *et al*., 2011). The phenotypes observed in our analysis during mitosis are consistent with published roles of Plk1 and the initital delay in localization of Rtnl1 at the spindle poles is due to the lack of astral microtubules present early in mitosis. Based on our injection of Binucleine 2 and BI25356, we conclude that Rtnl1 localization requires an active astral microtubule network.

## Discussion

Research over the last decade has highlighted the dramatic changes of the ER during cell division, however there is little known about the mechanistic properties that govern these mitotic changes (Lu *et al*., 2009; Bergman *et al*., 2015; Smyth *et al*., 2015). Our study has focused on the dynamics of the highly conserved ER shaping protein, Rtnl1, during mitosis. Here we show that Rtnl1 localizes at the spindle poles just prior to bulk of the ER membrane at prometaphase. Using precise temporal inhibition, we show that microtubule dynamics are necessary for proper ER localization and partitioning during mitosis. Furthermore, using small molecule inhibitors, Binucleine 2 and BI25356, that affect the formation of astral microtubule network early in mitosis leads to defects in ER localization at the spindle poles early in mitosis. This work highlights the mechanistic requirements of mitotic ER localization and provides a framework for ER partitioning during cell division.

Movement and changes of the ER during mitosis is a poorly understood topic, with several studies focused on the structural changes that occur during cell division (Puhka *et al*., 2007, 2012; Lu *et al*., 2009). It has been well established that organization of the ER relies on the microtubule network during interphase (Terasaki *et al*., 1986; Klopfenstein *et al*., 1998; Waterman-Storer and Salmon, 1998), however it is unknown if microtubules perform a similar role during mitosis. The ER shares a close localization with the mitotic spindle and poles and it has generally been assumed that microtubules and associated motor proteins are responsible for mitotic ER organization and partitioning. However, studies in *S. cerevisiae* and *C. elegans* have implicated the actin cytoskeleton network in mitotic ER dynamics (Poteryaev *et al*., 2005). Our data of the reliance of the mitotic spindle in mitotic ER organization is in line with recent studies that have demonstrated the requirement of centrosomes and astral microtubules in dividing Drosophila neuroblast (Smyth *et al*., 2015). Furthermore, recruitment of Rtnl1 to the spindle poles early in mitosis suggests the involvement of ER shaping proteins in the bulk movement of ER. Several studies have indicated a microtubule interaction between the mammalian homolog of Rtnl1 and the microtubule severing protein, Spastin (Mannan *et al*., 2006; Park *et al*., 2010) but how ER moves along the microtubule network is not known. A possibility is the kinesin-6 motor proteins, Mklp1, and Mklp2. These motor proteins are well noted for their role during mitosis, mainly cytokinesis (Gruneberg *et al*., 2004; Guse *et al*., 2005) and localize to the spindle midzone. Association of Rtnl1 with Mklp1 or Mklp2 would also explain the localization at the midbody during telophase (Fig. 1A, rightmost panel). In support of this association, Aurora B is well known for its regulation of Mklp1 and Mklp2 and Rtnl1 localization is absent from the midbody in our injections of Binucleine 2 during mitosis (Fig 4A, 12:30 time point).

## Materials and Methods

### Fly Stocks and Genetics

His2Av-RFP flies were a gift from Patrick O’Farrell (UCSF, San Francisco, CA). Rtnl1-GFP[G00199] stock was obtained from the Flytrap database. Lines were generated using protein-trap methodology (Morin *et al*., 2001). Lines were crossed, generating genotypes w;His2Av-RFP; P{PTT-GA} Pdi[G00198]/TM3. Homozygotes of Rtnl1[G00199]; His2Av-RFP or UAS-GFP-Lamin; UAS-RFP-KDEL were created by crossing individual fluorescent lines to balancer stocks and then mating the F1 progeny and driven using a tubulin-Gal4 line for expression in the early embryo.

### Embryo Collection and microinjection

Embryos were collected on grape-juice agar plates, aged on collection plates and dechorionated by hand. Dechorionated embryos were briefly desiccated and microinjected as previously described (Riggs *et al*., 2007).

Needle concentrations for injected solutions were as follows: Colchicine and Pacitaxel (5µM), cytochalasin D, Binucleine 2, and BI25356 (10µM). All drugs were dissolved in DMSO to a final concentration of 10%.

### Imaging live Embryos by Laser Confocal Microscopy

Confocal images of injected embryos were obtained on an inverted microscope (Zeiss Cell Observer, Carl Zeiss Microimaging, Inc.) using the 488nm and 543nm wavelengths from an Argon laser. Images were captured with a C-Apochromat 1.2 NA 100x objective (Carl Zeiss MicroImaging, Inc.) and analyzed with ImageJ (W. Rasband, National Institute of Health [NIH], Bethesda, MD) and Axiovision (Carl Zeiss MicroImaging, Inc).

## Supplemental figures

Supplemental Figure 1: **Stage 10 GFP-Lamin embryo imaged through mitosis.** GFP-Lamin localizes along the nuclear periphery at the start of mitosis. At 5:20 the nucleus starts to destabilize indicating the start of nuclear envelope breakdown (NEB), by 9:20 timepoint Lamin disappeared from the nucleus and reappeared at 15:00 at cytokinesis and nuclear envelope reformation.

