## Supplementary figures and images for "Microtubules are necessary for proper Reticulon localization during mitosis"

### Supplemental Table 1

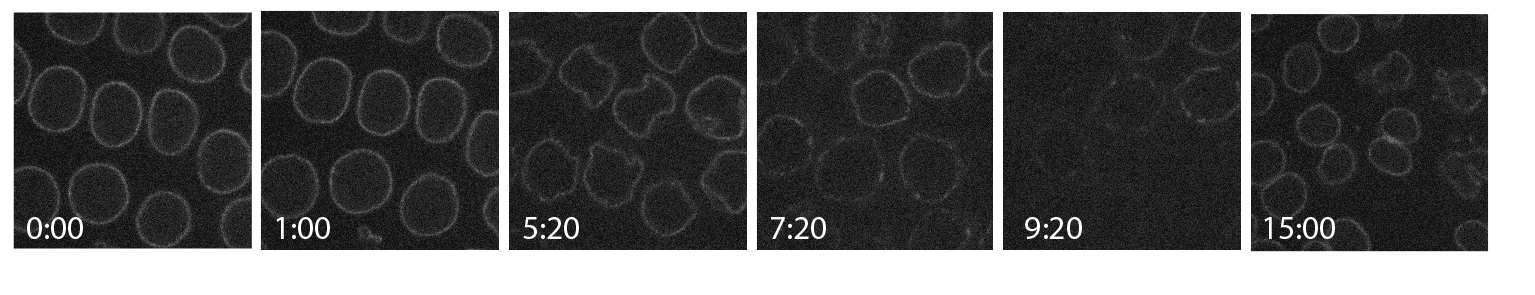
